# A Metabolomics Study of Thrombosis after Cardiac Surgery in Children with Congenital Heart Disease

**DOI:** 10.1101/2023.02.20.529324

**Authors:** Shengxu Li, Dave Watson, Alissa Jorgenson, Zainab Adelekan, Weihong Tang, Kathleen Garland, Leah Zupancich, David Dassenko, R. Erik Edens, David M. Overman, Marnie T. Huntley

## Abstract

**Objective:** Thrombosis is a major complication after cardiac surgery in children with congenital heart disease (CHD). The mechanisms underlying thrombosis development remain poorly understood. We aimed to identify novel circulating metabolites before cardiac surgery that are associated with the development of thrombosis after surgery in children with CHD.

**Approach and results:** All blood samples were drawn right before surgical incision and after the induction of anesthesia, and plasma was separated immediately under 4 °C. Untargeted metabolomic data were measured by Metabolon in plasma from children (age range: 0 days-18 years) with CHD undergoing cardiac surgery. The primary outcome was thrombosis within 30 days of surgery or before discharge. Associations of individual metabolites with thrombosis were assessed with logistic regression with false discovery rate (FDR) correction for multiple comparison and adjustment for clinical characteristics; elastic net regression was used to select a prediction model. Out of 1,115 metabolites measured in samples from 203 children, 776 met the quality control criteria. In total, 26 children (12.8%) developed thrombosis. Among the 776 metabolites, 195 were significantly associated with thrombosis (FDR q-value < 0.05). The top three metabolites showing the strongest associations with thrombosis were eicosapentaenoate, steroid monosulfate C19H28O6S, and formiminoglutamate (FDR = 0.01 for all). Pathway analysis showed that the pathways of nicotinate and nicotinamide metabolism (P = 0.0007) and glycerophospholipid metabolism (P = 0.002) were enriched and had significant impacts on the development of thrombosis. In elastic net regression analysis, the area under the receiver operating-characteristic curve of a prediction model for thrombosis was 0.94 in the training sample (70% of the total sample) and 0.84 in the testing sample (the remaining 30%).

**Conclusion:** We have identified promising novel metabolites and metabolic pathways associated with thrombosis. Future studies are warranted to confirm the findings and examine the mechanistic pathways to thrombosis.

## INTRODUCTION

Congenital heart disease (CHD) is the most common congenital malformation at birth, affecting about 1% of all live births in recent decades^1–3^. In the United States, approximately 40,000 children are born with CHD each year, with 25% requiring invasive intervention, usually surgery, within the first year of life^1^. Thrombosis is a common complication in children with CHD after corrective surgeries^4,5^. In a study from Canada, thrombi were observed in 11% of all surgeries in patients with CHD^6^. Another study including 91,909 children with CHD reported that children younger than 28 days had a high thrombosis rate of 61%^5^. Moreover, the incidence of thrombosis is likely underestimated due to a lack of systematic screening for thrombosis among CHD patients undergoing cardiac surgeries^7^. Once thrombosis has developed, only 62% of thrombotic events resolve^6^. Serious complications can develop in patients with thrombosis, which is associated with longer intensive care unit and hospital stays, higher odds of cardiac arrest, catheter reintervention, reoperation, and death^5,6^.

The mechanisms underlying the development of thrombosis after surgery in children with CHD are not well understood. It is known that hemostasis development begins in utero and evolves with age^8,9^. As a result, the still-developing hemostatic system has a limited capacity to maintain the delicate hemostatic balance^9^. In addition, many factors may contribute to the risk of thrombosis after surgery in children with CHD, including altered blood flow, endothelial disruption, exposure to prosthetic materials (especially central lines), inflammation and blood stream infection, and mechanical support^10^.

In recent years, metabolomic technology, which simultaneously measures thousands of metabolites in biofluids, has made it possible to detect metabolic alterations and to elucidate new metabolic mechanisms on a hypothesis-free basis^11^. Mass spectrometry coupled with liquid chromatography or gas chromatography—commonly referred to as the non-targeted metabolomic approach—has proved to be an exceedingly promising tool for studying complex pathophysiological processes^12^. For example, Wang et al. showed that the metabolite trimethylamine N-oxide (TMAO) was associated with future risk of atherosclerotic events, and they discovered a novel link between the gut-flora-dependent metabolism of dietary phosphatidylcholine and cardiovascular pathogenesis^13^. So far, few studies have used metabolomics technology to explore the etiology of thrombosis^14–18^, and no studies have used the platform to examine thrombosis after surgery in children with CHD. In this study, we aimed to identify novel metabolites and metabolic pathways that are associated with the development of thrombosis after cardiac surgery in children with CHD.

## METHODS

### Study sample

In this prospective study, children with CHD who planned to have corrective cardiac surgery at the Cardiovascular Care Center at Children’s Minnesota, the seventh largest stand-alone children’s hospital in the United States, were eligible for the study. Patients with the following conditions were excluded: 1) body weight < 1800 grams at the time of surgery; 2) receiving Extracorporeal Membrane Oxygenation at the time of surgery; 3) known clotting or bleeding disorder (e.g., Von Willebrand disease, Factor V Leiden, Factor VIII or Factor IX deficiency, methylenetetrahydrofolate reductase gene mutation); 4) isolated patent ductus arteriosus ligation; 5) parent or legal guardian unable to provide informed consent due to diminished capacity; and 6) prior enrollment in this study during the same hospitalization.

### Clinical data collection

Clinical data, including demographics, diagnoses, pre-operation treatments, surgical procedures, medical history (i.e., thrombosis or bleeding events), activated clotting time, and other relevant laboratory tests, were retrieved from the Electronic Medical Record (EMR) by A.J and Z.A. The primary outcome of thrombosis was determined by a thorough chart review of all EMRs for each patient stay. A thrombosis event was diagnosed by bedside vascular ultrasound, cardiac catheterization reports, or computed tomography angiography reports. Thrombus events included in this study were limited to those occurring within 30 days or before discharge (whichever was sooner). A final adjudication of all thrombi was performed by a physician (M.H.), who also reviewed all clinical data from 16 randomly selected patients. Clinical data reviewers (A.J., Z.A., and M.H) were blinded to metabolomic data.

### Blood sample collection and processing

Immediately after the induction of anesthesia and before incision, one milliliter of whole blood was drawn through a central line into an EDTA tube, which was then inverted 8-10 times to assure anticoagulation. The blood sample was immediately put into an ice bag and transferred to the lab at Children’s Minnesota for immediate separation of plasma. A laboratory technician manually further inverted the tube a minimum of 20 times prior to centrifuging the sample for 10-15 minutes at a minimum of 2000 g. Separated plasma was aliquoted into a pre-chilled, barcoded cryovial (0.7 mL) provided by Metabolon (Morrisville, NC, a service provider for the study). Plasma samples were flash-frozen at −80°C before being shipped to Metabolon for metabolomics quantification.

### Metabolomic quantification

Metabolon quantified all plasma samples, which were shipped in two separate batches. Metabolon used the untargeted, ultrahigh performance liquid chromatography-tandem mass spectroscopy-based metabolomic quantification protocol and followed rigorous protocols for sample handling, processing, and metabolites profiling, including the use of internal standards, blank samples, and technical replicates.

We performed additional quality assurance via 5 within-batch replicates and 10 between-batch replicates, which were blinded to Metabolon. Spearman correlations were calculated within all replicate groups for each metabolite. Only metabolites with call rates (i.e., non-missing data rates) ≥ 75% and with all correlations ≥0.5 were included for analysis. Among replicate pairs, one sample was randomly selected for analysis.

### Statistical analysis

#### Individual metabolite analysis

Logistic regression models were used to examine the associations of individual metabolites with the development of thrombosis after surgery, while adjusting for age, race, sex, personal history of thrombosis, personal history of postoperative bleeding, and STAT (i.e., Society of Thoracic Surgeons-European Association for Cardio-Thoracic Surgery) score. Concentrations of individual metabolites were batch-adjusted, median-imputed, (natural) log-transformed, and normalized to z-scores before further analysis, thus odds ratios (ORs) are interpretable for a one standard deviation change on the log scale. To control for multiple comparisons, the (positive) false discovery rate (FDR) was controlled at the 5% level, specifically using a 0.05 significance threshold for the q-value—the FDR analogue of the p-value^19,20^. ORs and 95% confidence intervals (CIs) were used to summarize the direction and magnitude of association.

#### Pathway analysis

We used MetaboAnalyst 5.0 (https://www.metaboanalyst.ca/MetaboAnalyst/home.xhtml), an online bioinformatics tool developed by Xia J., et al^21,22^, to analyze metabolomics pathways (a.k.a., MetPA)^23^. Metabolites that showed a significant association with thrombosis (i.e., q-value < 0.05) were input into the MetPA platform. In the current study, pathway enrichment was determined by over-representation analysis, and the pathway impact was determined by the sum of the importance measures of the matched metabolites normalized by the sum of the importance measures of all metabolites in each pathway.

#### Prediction model

Elastic net logistic regression was used to develop a predictive model for thrombosis^24^. The sample was randomly split into two sets: a training set that included 70% of patients and a test set with the remaining 30%. Within the training set, 10-fold cross validation was used to select the parameters alpha and lambda (i.e., the elastic net mixing and penalty parameters, respectively)^25^. Alpha was chosen to minimize the deviance measure, and lambda was chosen to be one standard error away from the minimizing lambda. Three separate sets of predictors were considered: only clinical information, only metabolites, and both clinical and metabolite information. Area under the receiver operating-characteristic curve (AUC) was used to evaluate and compare the performance of prediction models in both training and test sets^26^.

## RESULTS

From December 2019 to July 2021, 203 children aged 0-18 years were enrolled, with 6 patients having two surgical encounters. A total of 235 plasma samples were quantified by Metabolon. Among the 1,115 metabolites that were common to the two batches, 877 metabolites had a call rate of at least 75%; among the 877 metabolites, 776 that had a Spearman correlation coefficient of at least 0.5 (for both between- and within-batch replicates) were included for final analysis (**Supplementary Figure 1**). For the 6 patients who were enrolled twice, data from the first of their two encounters were used. Among the 203 patients, the median age was 1.4 (interquartile range: 0.4-6.9) years, 87 (42.9%) were girls, and 75.9% were white, 7.4% Black, and 16.7% other races (**Table 1**). Other clinical characteristics of the cohort are shown in **Table 1**.

**Table 1.** Characteristics of the study sample before surgery (n=203)

| Variable | Mean±SD, median (interquartile range),<br>or n (%) |
| --- | --- |
| Age (year) | 1.4 (0.4-6.9) |
| Female, n (%) | 87 (42.9) |
| Race, n (%) |  |
| White | 154 (75.9) |
| Black | 15 (7.4) |
| Other | 34 (16.7) |
| Medical history of bleeding, n (%) | 9 (4.4) |
| Medical history of thrombosis | 31 (15.3) |
| STAT score, n (%) |  |
| 1 | 70 (34.5) |
| 2-3 | 94 (46.3) |
| 4-5 | 39 (19.2) |
| Anticoagulant medication, n (%) | 55 (27.1) |

Twenty-six patients (12.8%) developed thrombosis after surgery. Among clinical characteristics, age was inversely associated with risk of thrombosis, and medical history of bleeding and thrombosis, STAT score, and pre-operation anti-coagulation medication were positively associated with risk of thrombosis after surgery (**Table 2**).

**Table 2.** Associations of potential risk factors with thrombosis

| Variable | Odds ratio | 95% confidence interval |  | p |
| --- | --- | --- | --- | --- |
|  |  | Lower limit | Upper limit |  |
| Age (year) | 0.73 | 0.58 | 0.92 | 0.009 |
| Female | 1.99 | 0.86 | 4.58 | 0.11 |
| Race (reference: white) |  |  |  |  |
| Black | 1.58 | 0.41 | 6.08 | 0.50 |
| Other | 0.40 | 0.09 | 1.78 | 0.23 |
| Medical history of bleeding | 6.25 | 1.56 | 25.05 | 0.01 |
| Medical history of thrombosis | 4.64 | 1.87 | 11.56 | 0.001 |
| STAT score (reference: 1) |  |  |  |  |
| 2-3 | 9.14 | 1.15 | 72.60 | 0.04 |
| 4-5 | 38.64 | 4.83 | 309.20 | 0.0006 |
| Anti-coagulation medication | 3.21 | 1.38 | 7.47 | 0.007 |

All variables were included in one model.

In individual metabolite analysis, 195 metabolites were associated with thrombosis after surgery with a q-value <0.05 (**Supplementary Table 1**). The number of significant metabolites was substantially more than expected by chance alone (39, see **Supplementary Figure 2** for quantile-quantile and volcano plots). Among the significant metabolites, 78 showed an inverse association and 117 a positive association with risk of thrombosis after surgery. The three metabolites with the lowest p-values were eicosapentaenoate (EPA) (OR=3.12; 95% CI=1.76-5.52), andro steroid monosulfate C19H28O6S (OR=4.07; 95% CI=1.99-8.36), and formiminoglutamate (OR=3.75; 95% CI=1.86-7.55). Of note, isoleucine (one of the top 20 significant metabolites; OR=2.29 and 95% CI=1.41-3.71), along with leucine (OR=2.25; 95% CI=1.39-3.65) and valine (OR=2.75; 95% CI=1.29-5.85), was associated with increased risk of developing thrombosis (**Supplementary Table 1**). In addition, the levels of acetylcarnitine, cerotoylcarnitine, lignoceroylcarnitine, 3-hydroxyoctanoylcarnitine, and carnitine in our study were associated with reduced risk of thrombosis (**Supplementary Table 1)**.

All 195 significant metabolites were input into MetaboAnalyst 5.0. Four pathways were significantly enriched at an FDR of 0.05 (**Table 3**); of these pathways, two also showed a large pathway impact on risk of thrombosis: glycerophospholipid metabolism and nicotinate and nicotinamide metabolism (**Figure 1**). In the glycerophospholipid metabolism pathway, the 7 significant metabolites were ethanolamine phosphate, phosphatidylethanolamine, sn-glycero-3-phosphoethanolamine, sn-glycerol-3-phosphate, choline phosphate, phosphatidyl choline, and 1-acyl-sn-glycero-3-phosphocholine. In the nicotinate and nicotinamide metabolism pathway, L-aspartate, quinolinate, nicotinamide, 1-methylnicotinamide, and n1-methyl-2-pyridone-5-carboxamine) were significant.

**Table 3.** Significant pathways identified via MetaboAnalyst 5.0 using 195 statistically significant metabolites from individual metabolite analysis

| Pathway | Number of metabolites in the pathway | Expected number of metabolites under the null | Observed number of metabolites | P value | FDR q value | Pathway impact |
| --- | --- | --- | --- | --- | --- | --- |
| Aminoacyl-tRNA biosynthesis | 48 | 1.73 | 10 | 3.79E-06 | 3.19E-4 | 0 |
| Valine, leucine and isoleucine biosynthesis | 8 | 0.29 | 4 | 9.62E-05 | 3.28E-3 | 0 |
| Nicotinate and nicotinamide metabolism | 15 | 0.54 | 5 | 1.17E-4 | 3.28E-3 | 0.33 |
| Glycerophospholipid metabolism | 36 | 1.30 | 7 | 2.03E-4 | 4.26E-3 | 0.37 |

FDR: false discovery rate

**Figure 1.**
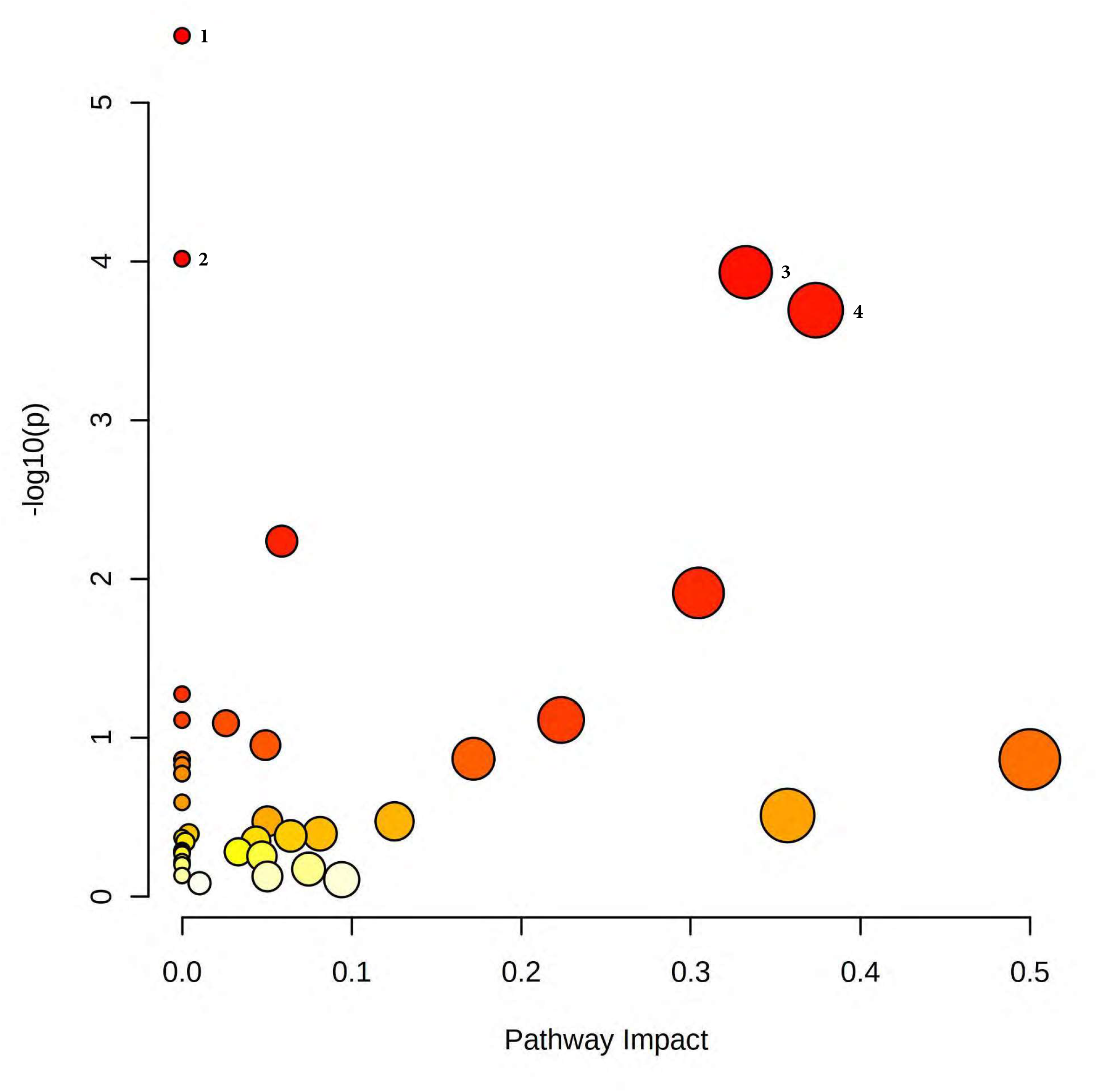
Pathway analysis results from MetaboAnalyst 5.0 based on 195 significant metabolites. 1. Aminoacyl-tRNA biosynthesis; 2. Valine, leucine and isoleucine biosynthesis; 3. Nicotinate and nicotinamide metabolism; and 4. Glycerophospholipid metabolism

In the final analysis, we used elastic net regression to develop a prediction model for thrombosis after surgery. The elastic net regression model with only clinical data selected age, sex, history of thrombosis, and STAT score as predictors (**Supplementary Table 2)**, and the AUC was 0.895 in the training set and 0.786 in the testing set (**Figure 2**). In the metabolites-only model, the elastic net regression selected 46 metabolites (**Supplementary Table 2**), and the AUC was 0.887 in the training set and 0.854 in the testing set (**Figure 2**). Finally, the model that included both clinical data and metabolites selected history of thrombosis, STAT score, and 17 metabolites (**Supplementary Table 2)**, and the AUC was 0.942 in the training set and 0.839 in the testing set (**Figure 2**). Compared to the clinical data-only model, the differences between AUCs of 0.053 for the metabolites-only model and 0.069 for the model with both clinical data and metabolites were not statistically significant (p=0.64 and p=0.59, respectively). The predicted probabilities for thrombosis development according to these models are shown in **Figure 3**. According to the model that included both clinical variables and metabolites, the predicted probability for developing thrombosis among those who developed thrombosis was > 12.5% for all but one in the testing set (i.e., 6/7).

**Figure 2.**
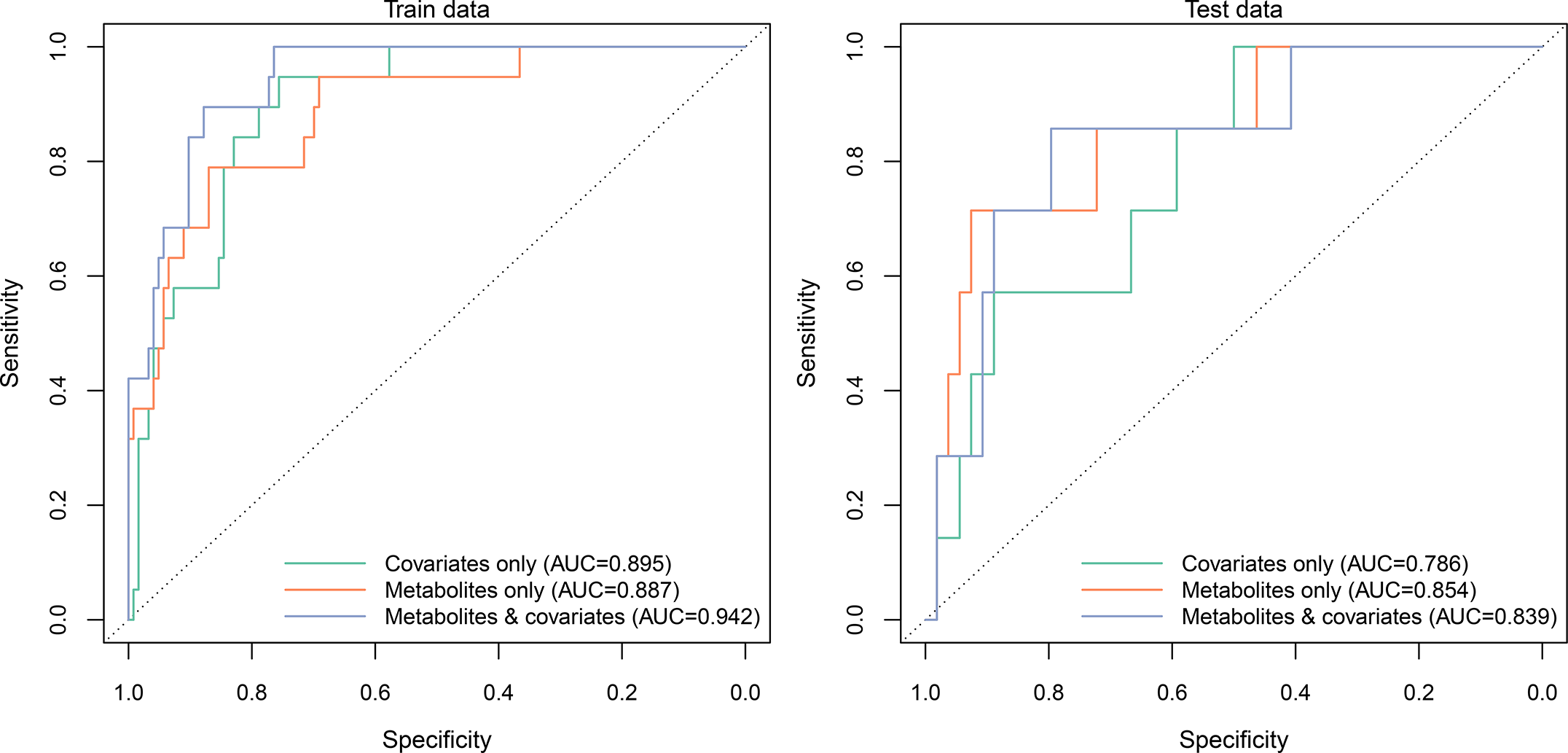
The area under the receiver operating characteristic curve of predictive models in the training sample (left) and in the testing sample (right)

**Figure 3.**
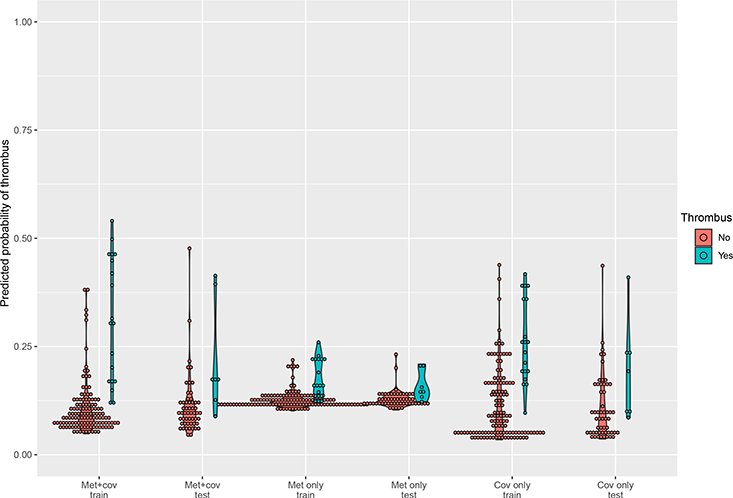
Predicted probability for developing thrombosis after surgery in children with congenital heart disease in the training sample and in the testing sample

## DISCUSSION

The current study identified novel individual metabolites and metabolic pathways that are associated with thrombosis after cardiac surgery in children with CHD. Prediction models with metabolites had better (though not statistically significantly better) performance with respect to AUC, suggesting the future utility of such models in clinical care. The results suggest that metabolomic technology is a powerful tool for studying the development of thrombosis in this vulnerable pediatric population.

Thrombosis after cardiac surgery in children with CHD is common, yet the underlying mechanisms for its development are not fully understood. Known contributing factors include immature hemostasis in children younger than 16 years, surgery trauma, line exposure, hypoxia, and infection. Although we identified 195 significant metabolites future work needs to confirm the findings of the current study and to start to understand what role these metabolites play in the development of thrombosis.

Limited literature is available for comparison with the findings of the current study. Among the top 20 metabolites, only EPA and isoleucine have been reported to be associated with thrombosis^18^. However, contrary to an earlier report that EPA is linked to reduced risk of thrombosis, EPA was associated with increased risk of thrombosis in the current study. The reason for this discrepancy is unclear. Confounding might have played a role, which is supported by the lack of benefit for an EPA supplement in a randomized clinical trial^27^. Associations of the three branched-chain amino acids (isoleucine, leucine, and valine) with increased risk of developing thrombosis are consistent with the fact that their catabolism promotes thrombosis risk by enhancing tropomodulin-3 propionylation in platelets^28^. For acylcarnitines, their levels are lower in patients after venous thromboembolism than in controls, and acylcarnitines are anticoagulants that inhibit factor Xa^14^.

The identification of novel metabolites for thrombosis is instrumental to finding new therapeutic or intervention measures. As an example, after TMAO’s role in thrombosis was confirmed^15^, the underlying mechanisms for its role were examined^29^, and targeting TMAO has become a new therapeutic strategy^30^. Although significant work remains to be done to translate the identification of novel metabolites in the current study to the development of new strategies targeting these metabolites and their metabolic pathways, our study demonstrates the feasibility and utility of the metabolomics approach in tackling the challenges in understanding thrombosis development after surgery in children with CHD.

Although the vast majority of metabolites identified in the current study do not have clear relevance to thrombosis, these identified metabolites do point to novel metabolic pathways. The glycerophospholipid metabolism pathway and the nicotinate and nicotinamide metabolism pathway had significant pathway impact on the development of thrombosis. Phosphoinositide 3-kinases, key players in the pathway, are known to be critical to various aspects of platelet biology^31^. Anti-phospholipid antibodies have been shown to increase the risk of thrombosis^32,33^. It has been reported that deficiency of coline, a key metabolite in the pathway, is associated with the risk of catheter-related venous thrombosis^34^. Taken together, these findings suggest the relevance of the glycerophospholipid metabolism pathway to the development of thrombosis. The nicotinate and nicotinamide metabolism pathway plays a key role in redox metabolism^35^. Redox metabolism is known to be linked to the development of thrombosis, which is partly mediated by platelet activation promoted by the nicotinamide adenine dinucleotide phosphate (NADPH) oxidase, a key target of oxidative stress-mediated platelet activation and thrombosis^36^. However, a full depiction of how the pathway is linked to the risk of thrombosis is lacking. Future studies are needed to investigate the underlying mechanisms by which the nicotinate and nicotinamide metabolism pathway plays a role in thrombosis development.

The prediction models we developed in the current study performed reasonably well in the testing sample. Although we observed increases of more than 5 percentage points in absolute AUC values for models including metabolomic data compared to the model with only clinical data, the differences were not significant, most likely because the sample size and the number of thrombosis events in the testing set were small. However, these models do suggest the potential utility of using a small set of metabolites in predicting risk of thrombosis after surgery in children with CHD, which should be further confirmed in future studies.

The current study has several strengths. It is a prospective study with rigorous quality assurance/control protocols. We collected the blood specimens before surgery, which ensured that the metabolomic profiles preceded the development of thrombosis events. The consistency in the timing of blood collection (i.e., right before incision) and in specimen handling (i.e., transfer on ice and immediate processing) ensured homogeneous conditions and minimized non-physiological concentration variations in the metabolites quantified. Both within- and between-batch blind duplicates provided critical information for data quality in the metabolomic quantification process. All these measures ensured high quality data on metabolite levels.. Finally, we demonstrated that elastic regression analysis is a powerful tool for selecting biological markers for clinical outcomes, even when the biological markers significantly outnumber the samples. The current study also had limitations. First, the sample size and the number of events were small. However, we were still able to identify numerous metabolites robustly associated with the development of thrombosis after taking multiple comparisons into account. Second, we did not include some important lab values related to thrombosis (e.g., international normalized ratio) in our prediction models because missing data would have reduced the already small sample size. These lab measurements showed modest association with thrombosis and future work might consider their association with metabolomic data. Finally, the patient population was children with CHD who needed corrective surgery, which limits the generalizability of our findings to other populations. Nevertheless, we believe that some of our findings may be applicable to other populations, which should be confirmed in future studies.

In conclusion, we have identified novel metabolites associated with the development of thrombosis after cardiac surgery in children with CHD. The new metabolism pathways indicated in the current study point to new directions for research aiming to elucidate the pathophysiological mechanisms underlying the development of thrombosis.

## ACKNOWLEDGEMENTS

The current study was supported by the Children’s Minnesota Research Committee and Children’s Minnesota Foundation. The funding agency did not have any role in conducting the study or in interpreting the final results.

The current study involved many colleagues at Children’s Minnesota, including colleagues at the Minneapolis Central Lab, the Cardiovascular Care Center, the Children’s Heart Clinic, and anesthesiologists, whose contributions are greatly appreciated. The authors of the study also wish to thank the children who participated in the study and their families. The authors also wish to thank Martin Cozza for his help with editing the manuscript.

## Figure legends

**Supplementary Figure 1**. Flow chart for filtering metabolites according to data quality measures

**Supplementary Figure 2**. For associations of individual metabolites with thrombosis, Quantile-quantile plot of observed versus expected p-values and volcano plot of p-values against effect sizes (i.e., odds ratios)

