## Supplementary material for "A Metabolomics Study of Thrombosis after Cardiac Surgery in Children with Congenital Heart Disease"

Supplementary Figure 1.

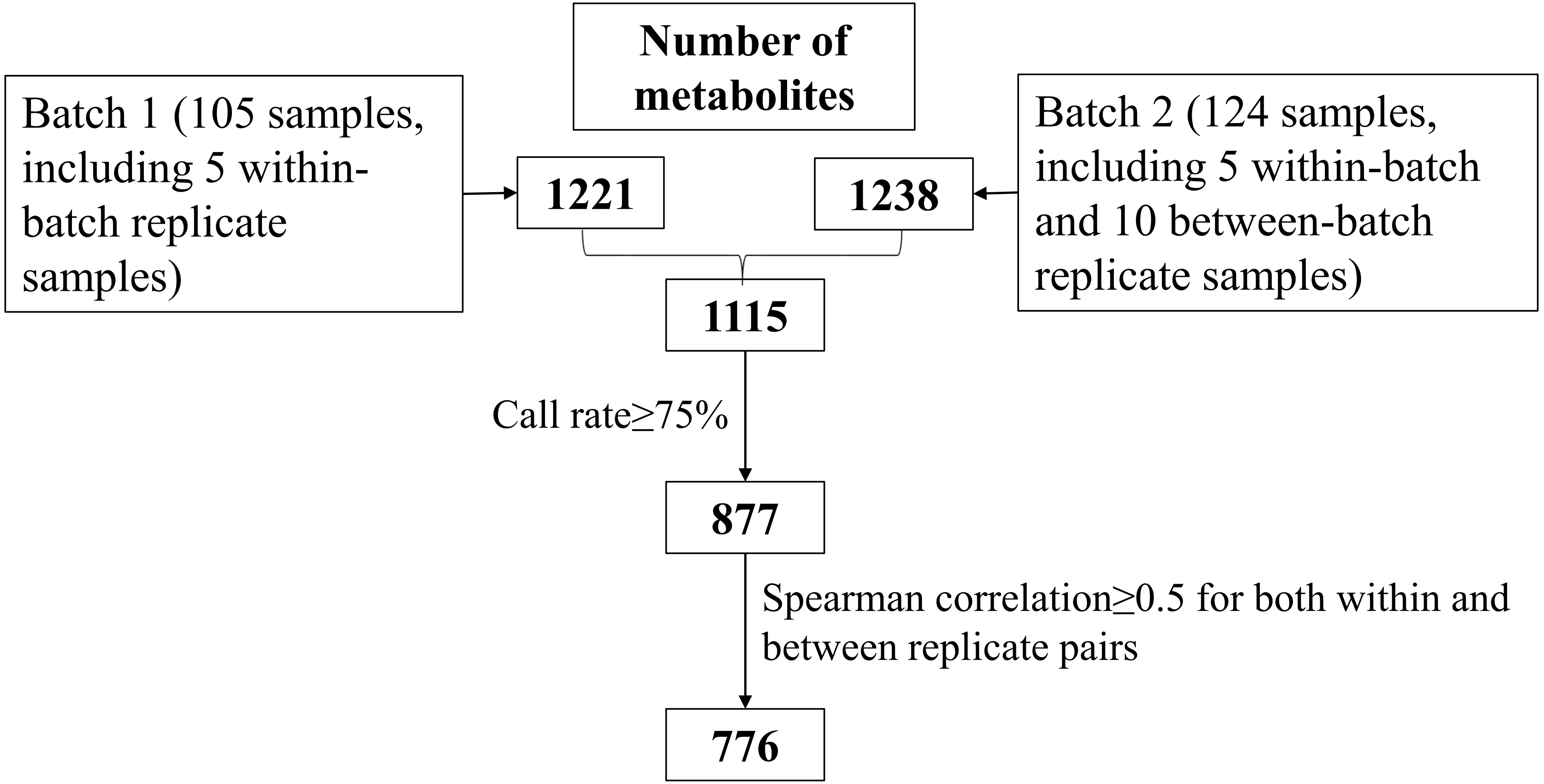

**Supplementary Table 1. Significant metabolites with a false discovery rate q-value less than 0.05**

| Chemical Name | Human Metabolome Database (HMDB) ID | Odds Ratio and 95% CI | p value | q value |
|---|---|---|---|---|
| eicosapentaenoate (EPA; 20:5n3) | HMDB0001999 | 3.12 (1.76, 5.52) | 0.0001 | 0.010473434 |
| andro steroid monosulfate C19H28O6S (1)* | HMDB0002759 | 4.07 (1.99, 8.36) | 0.0001 | 0.010473434 |
| formiminoglutamate | HMDB0000854 | 3.75 (1.86, 7.55) | 0.0002 | 0.010473434 |
| stearidonate (18:4n3) | HMDB0006547 | 2.97 (1.64, 5.37) | 0.0003 | 0.010473434 |
| androstenediol (3beta,17beta) disulfate (2) | HMDB0240313 | 3.38 (1.74, 6.58) | 0.0003 | 0.010473434 |
| gamma-glutamylleucine | HMDB0011171 | 2.65 (1.55, 4.50) | 0.0003 | 0.010473434 |
| N-acetylkynurenine (2) | HMDB0240342 | 3.16 (1.68, 5.95) | 0.0004 | 0.010473434 |
| kynurenate | HMDB0000715 | 3.37 (1.72, 6.62) | 0.0004 | 0.010473434 |
| 2-linoleoylglycerol (18:2) | HMDB0011538 | 3.51 (1.74, 7.07) | 0.0004 | 0.010473434 |
| 1-palmitoleoylglycerol (16:1)* | HMDB0011565 | 4.10 (1.86, 9.05) | 0.0005 | 0.010473434 |
| 1-oleoylglycerol (18:1) | HMDB0011567 | 3.33 (1.69, 6.58) | 0.0005 | 0.010473434 |
| 2R,3R-dihydroxybutyrate | HMDB0000498 | 3.22 (1.65, 6.28) | 0.0006 | 0.010473434 |
| X-25433 | | 5.79 (2.13, 15.76) | 0.0006 | 0.010473434 |
| X-24544 | | 3.85 (1.76, 8.39) | 0.0007 | 0.010473434 |
| gamma-glutamylphenylalanine | HMDB0000594 | 2.49 (1.47, 4.21) | 0.0007 | 0.010473434 |
| N2,N2-dimethylguanosine | HMDB0004824 | 3.65 (1.72, 7.75) | 0.0008 | 0.010473434 |
| isoleucine | HMDB0000172 | 2.29 (1.41, 3.71) | 0.0008 | 0.010473434 |
| X-21410 | | 4.36 (1.85, 10.28) | 0.0008 | 0.010473434 |
| 16alpha-hydroxy DHEA 3-sulfate | HMDB0062544 | 3.78 (1.73, 8.27) | 0.0009 | 0.010473434 |
| androstenediol (3beta,17beta) disulfate (1) | HMDB0240313 | 2.91 (1.55, 5.46) | 0.0009 | 0.010473434 |
| gamma-glutamylisoleucine* | HMDB0011170 | 2.37 (1.42, 3.94) | 0.0009 | 0.010473434 |
| hydroxy-CMPF* | | 3.16 (1.60, 6.24) | 0.0009 | 0.010473434 |
| N6-carbamoylthreonyladenosine | HMDB0041623 | 3.47 (1.66, 7.23) | 0.0009 | 0.010473434 |
| 1-arachidonylglycerol (20:4) | HMDB11578 | 2.93 (1.55, 5.54) | 0.0009 | 0.010473434 |
| leucine | HMDB0000687 | 2.25 (1.39, 3.65) | 0.0010 | 0.010473434 |
| 2'-deoxyuridine | HMDB0000012 | 3.26 (1.61, 6.59) | 0.0010 | 0.010473434 |
| docosahexaenoate (DHA; 22:6n3) | HMDB0002183 | 3.97 (1.74, 9.05) | 0.0011 | 0.010473434 |
| 3-carboxy-4-methyl-5-propyl-2-furanpropanoate (CMPF) | HMDB0061112 | 2.89 (1.53, 5.46) | 0.0011 | 0.010473434 |
| X-24328 | | 3.02 (1.56, 5.88) | 0.0011 | 0.010473434 |
| taurocholate | HMDB0000036 | 3.28 (1.61, 6.68) | 0.0011 | 0.010473434 |
| N-acetylvaline | HMDB0011757 | 3.32 (1.61, 6.83) | 0.0011 | 0.010473434 |
| 1-linoleoylglycerol (18:2) | HMDB0011568 | 2.90 (1.52, 5.52) | 0.0012 | 0.01053218 |
| phosphoethanolamine | HMDB0000224 | 2.73 (1.49, 5.03) | 0.0012 | 0.01053218 |
| N-acetylisoleucine | HMDB0061684 | 2.68 (1.47, 4.87) | 0.0012 | 0.01053218 |
| 1-methylnicotinamide | HMDB0000699 | 2.99 (1.53, 5.82) | 0.0013 | 0.010761781 |
| N-acetylphenylalanine | HMDB0000512 | 2.73 (1.48, 5.03) | 0.0013 | 0.010761781 |
| pregnenolone sulfate | HMDB0000774 | 3.43 (1.61, 7.32) | 0.0014 | 0.011147018 |
| bilirubin degradation product, C16H18N2O5 (3)** | | 2.55 (1.43, 4.53) | 0.0014 | 0.011147018 |
| caprylate (8:0) | HMDB0000482 | 2.32 (1.38, 3.90) | 0.0015 | 0.011147018 |
| 21-hydroxypregnenolone disulfate | | 2.93 (1.50, 5.73) | 0.0017 | 0.011692506 |
| 2-stearoyl-GPE (18:0)* | HMDB0011129 | 2.81 (1.47, 5.36) | 0.0017 | 0.011692506 |
| 5-methyluridine (ribothymidine) | HMDB0000884 | 3.04 (1.52, 6.08) | 0.0017 | 0.011692506 |
| N-acetylalanine | HMDB0000766 | 3.48 (1.59, 7.60) | 0.0018 | 0.011692506 |
| gamma-glutamylthreonine | HMDB0029159 | 2.53 (1.41, 4.52) | 0.0018 | 0.011692506 |
| X-15503 | | 2.47 (1.39, 4.37) | 0.0019 | 0.012681886 |
| picolinoylglycine | HMDB0059766 | 2.19 (1.33, 3.59) | 0.0021 | 0.013073502 |

| | | | | |
|---|---|---|---|---|
| N-acetyltyrosine | HMDB0000866 | 1.98 (1.28, 3.05) | 0.0021 | 0.013073502 |
| C-glycosyltryptophan | HMDB0240296 | 2.97 (1.48, 5.97) | 0.0022 | 0.013182977 |
| 1-palmitoleoyl-GPC (16:1)* | HMDB0010383 | 2.13 (1.31, 3.47) | 0.0022 | 0.013182977 |
| glycerol 3-phosphate | HMDB0000126 | 2.64 (1.41, 4.92) | 0.0023 | 0.013182977 |
| taurochenodeoxycholate | HMDB0000951 | 3.34 (1.54, 7.26) | 0.0023 | 0.013182977 |
| 7-methylguanine | HMDB0000897 | 3.71 (1.59, 8.63) | 0.0024 | 0.013182977 |
| gluconate | HMDB0000625 | 1.91 (1.26, 2.89) | 0.0024 | 0.013182977 |
| 1-(1-enyl-palmitoyl)-2-linoleoyl-GPC (P-16:0/18:2)* | HMDB0011211 | 0.37 (0.19, 0.70) | 0.0024 | 0.013182977 |
| (N(1) + N(8))-acetylspermidine | HMDB0002189,HMDB0001276 | 3.31 (1.52, 7.22) | 0.0026 | 0.014017867 |
| X-13866 | | 2.12 (1.30, 3.47) | 0.0027 | 0.014109965 |
| bilirubin degradation product, C16H18N2O5 (2)** | | 2.42 (1.36, 4.30) | 0.0027 | 0.014109965 |
| docosapentaenoate (n3 DPA; 22:5n3) | HMDB0006528,HMDB0001976 | 2.17 (1.30, 3.63) | 0.0030 | 0.014832481 |
| docosapentaenoate (n6 DPA; 22:5n6) | HMDB0001976 | 2.27 (1.32, 3.91) | 0.0030 | 0.014832481 |
| N-acetylhistidine | HMDB0032055 | 3.10 (1.47, 6.54) | 0.0030 | 0.014832481 |
| caprate (10:0) | HMDB0000511 | 2.33 (1.33, 4.07) | 0.0031 | 0.014847733 |
| cortisone | HMDB0002802 | 4.08 (1.60, 10.40) | 0.0032 | 0.01513145 |
| kynurenine | HMDB0000684 | 2.82 (1.41, 5.61) | 0.0033 | 0.01513145 |
| 1-arachidonoyl-GPI (20:4)* | HMDB0061690 | 2.39 (1.33, 4.28) | 0.0034 | 0.015422473 |
| X-21364 | | 2.42 (1.34, 4.39) | 0.0035 | 0.015701888 |
| dehydroepiandrosterone sulfate (DHEA-S) | HMDB0001032 | 3.22 (1.47, 7.09) | 0.0036 | 0.015975557 |
| bilirubin degradation product, C17H18N2O4 (2)** | | 2.24 (1.30, 3.86) | 0.0037 | 0.016064192 |
| biliverdin | HMDB0001008 | 2.46 (1.34, 4.51) | 0.0038 | 0.016064192 |
| gamma-glutamylmethionine | HMDB0029155 | 2.14 (1.28, 3.57) | 0.0038 | 0.016064192 |
| glycerophosphoethanolamine | HMDB0000114 | 2.39 (1.32, 4.33) | 0.0040 | 0.016322439 |
| xanthurenate | HMDB0000881 | 2.19 (1.28, 3.73) | 0.0040 | 0.016322439 |
| nicotinamide | HMDB0001406 | 2.43 (1.33, 4.45) | 0.0040 | 0.016322439 |
| deoxycholic acid glucuronide | | 0.44 (0.25, 0.77) | 0.0041 | 0.016322439 |
| bilirubin degradation product, C17H18N2O4 (3)** | | 2.27 (1.29, 3.99) | 0.0043 | 0.016803018 |
| 3-methylcytidine | HMDB0240577 | 3.01 (1.41, 6.42) | 0.0043 | 0.016803018 |
| hydroxyasparagine** | HMDB32332 | 2.35 (1.31, 4.22) | 0.0044 | 0.016803018 |
| 3-(3-amino-3-carboxypropyl)uridine* | | 2.91 (1.39, 6.09) | 0.0045 | 0.017060993 |
| 1-linoleoyl-GPG (18:2)* | HMDB0240600 | 2.61 (1.35, 5.07) | 0.0045 | 0.017083702 |
| choline phosphate | HMDB0001565 | 2.43 (1.31, 4.51) | 0.0049 | 0.018068048 |
| 1-stearoyl-GPE (18:0) | HMDB0011130 | 2.30 (1.28, 4.12) | 0.0054 | 0.01919833 |
| threonine | HMDB0000167 | 2.20 (1.26, 3.83) | 0.0054 | 0.01919833 |
| pelargonate (9:0) | HMDB0000847 | 2.10 (1.24, 3.54) | 0.0054 | 0.01919833 |
| X-17010 | | 2.60 (1.33, 5.11) | 0.0054 | 0.01919833 |
| mannonate* | | 1.79 (1.19, 2.71) | 0.0056 | 0.019250003 |
| N1-methylinosine | HMDB0002721 | 2.95 (1.37, 6.33) | 0.0056 | 0.019250003 |
| methionine sulfoxide | HMDB0002005 | 1.88 (1.20, 2.93) | 0.0057 | 0.019250003 |
| 5alpha-pregnan-3beta,20alpha-diol disulfate | HMDB0094650 | 1.98 (1.22, 3.20) | 0.0057 | 0.019250003 |
| gamma-glutamyl-alpha-lysine | | 2.14 (1.25, 3.69) | 0.0059 | 0.019552651 |
| taurocholenate sulfate* | | 3.04 (1.37, 6.73) | 0.0061 | 0.020122272 |
| hydroxy-N6,N6,N6-trimethyllysine* | | 2.15 (1.24, 3.71) | 0.0063 | 0.020398202 |
| N-acetylputrescine | HMDB0002064 | 3.46 (1.42, 8.45) | 0.0064 | 0.020398202 |
| 1-palmitoyl-2-oleoyl-GPE (16:0/18:1) | HMDB0005320 | 2.14 (1.24, 3.70) | 0.0064 | 0.020398202 |
| picolinate | HMDB0002243 | 2.15 (1.24, 3.74) | 0.0066 | 0.02067051 |
| alanine | HMDB0000161 | 2.38 (1.27, 4.45) | 0.0066 | 0.02067051 |

| | | | | |
|---|---|---|---|---|
| sphingosine | HMDB0000252 | 2.44 (1.28, 4.65) | 0.0070 | 0.021227357 |
| betaine | HMDB0000043 | 0.44 (0.24, 0.80) | 0.0072 | 0.021227357 |
| bilirubin degradation product, C17H18N2O4 (1)** | | 2.10 (1.22, 3.61) | 0.0073 | 0.021227357 |
| N-formylphenylalanine | HMDB0240317 | 2.22 (1.24, 3.97) | 0.0073 | 0.021227357 |
| methionine | HMDB0000696 | 1.78 (1.17, 2.72) | 0.0074 | 0.021227357 |
| N-acetyltryptophan | HMDB0013713 | 1.94 (1.19, 3.14) | 0.0075 | 0.021227357 |
| linoleoyl-arachidonoyl-glycerol (18:2/20:4) [2]* | HMDB0007257 | 2.31 (1.25, 4.27) | 0.0075 | 0.021227357 |
| X-12015 | | 2.47 (1.27, 4.81) | 0.0076 | 0.021227357 |
| 1-stearoyl-2-oleoyl-GPE (18:0/18:1) | HMDB0008993 | 2.08 (1.22, 3.57) | 0.0076 | 0.021227357 |
| beta-citrylglutamate | | 3.18 (1.36, 7.44) | 0.0076 | 0.021227357 |
| N-acetylglucosamine/N-acetylgalactosamine | HMDB0000212,HMDB0000215 | 2.38 (1.26, 4.51) | 0.0076 | 0.021227357 |
| X-11470 | | 0.48 (0.28, 0.82) | 0.0077 | 0.021227357 |
| 1-stearoyl-2-oleoyl-GPC (18:0/18:1) | HMDB0008038 | 2.20 (1.23, 3.92) | 0.0079 | 0.021371003 |
| 2-piperidinone | HMDB0011749 | 0.46 (0.26, 0.82) | 0.0079 | 0.021371003 |
| 2,3-dihydroxy-5-methylthio-4-pentenoate (DMTPA)* | HMDB0240388 | 2.10 (1.21, 3.64) | 0.0080 | 0.021371003 |
| 1-arachidonoyl-GPE (20:4n6)* | HMDB0011517 | 1.90 (1.18, 3.05) | 0.0081 | 0.021621137 |
| sphingomyelin (d18:1/18:1, d18:2/18:0) | HMDB0012101 | 2.10 (1.21, 3.63) | 0.0084 | 0.022253049 |
| 5-hydroxylysine | HMDB0000450 | 3.28 (1.35, 7.95) | 0.0087 | 0.022692567 |
| valine | HMDB0000883 | 2.75 (1.29, 5.85) | 0.0088 | 0.022692567 |
| gamma-glutamylvaline | HMDB0011172 | 2.10 (1.20, 3.67) | 0.0092 | 0.023628929 |
| ribitol | HMDB0002917,HMDB0001851,HMDB0000568,HMDB0000508 | 1.93 (1.18, 3.18) | 0.0094 | 0.023960979 |
| glycocholate | HMDB0000138 | 2.20 (1.21, 4.02) | 0.0100 | 0.025172116 |
| 1-palmitoyl-2-oleoyl-GPC (16:0/18:1) | HMDB0007972 | 2.04 (1.18, 3.51) | 0.0103 | 0.02546266 |
| 1-palmitoyl-GPE (16:0) | HMDB0011503 | 2.13 (1.19, 3.81) | 0.0104 | 0.02546266 |
| erythronate* | HMDB0000613 | 2.28 (1.21, 4.28) | 0.0105 | 0.02546266 |
| N1-methyladenosine | HMDB0003331 | 2.86 (1.28, 6.40) | 0.0105 | 0.02546266 |
| aspartate | HMDB0000191 | 1.84 (1.15, 2.94) | 0.0106 | 0.02546266 |
| 5-methylthioadenosine (MTA) | HMDB0001173 | 2.01 (1.18, 3.44) | 0.0106 | 0.02546266 |
| N-acetylserine | HMDB0002931 | 2.75 (1.26, 5.99) | 0.0108 | 0.025601964 |
| pseudouridine | HMDB0000767 | 2.46 (1.23, 4.93) | 0.0111 | 0.026121904 |
| 1-oleoyl-GPC (18:1) | HMDB0002815 | 1.89 (1.16, 3.09) | 0.0112 | 0.026121904 |
| X-21471 | | 2.77 (1.26, 6.08) | 0.0113 | 0.026121904 |
| N-acetylarginine | HMDB0004620 | 2.14 (1.19, 3.85) | 0.0114 | 0.026121904 |
| 1-arachidonoyl-GPC (20:4n6)* | HMDB0010395 | 1.86 (1.15, 3.01) | 0.0114 | 0.026121904 |
| X-21816 | | 1.94 (1.16, 3.26) | 0.0120 | 0.027173893 |
| acetylcarnitine (C2) | HMDB0000201 | 0.52 (0.31, 0.87) | 0.0121 | 0.027173893 |
| 2-O-methylascorbic acid | HMDB0240294 | 2.06 (1.17, 3.63) | 0.0122 | 0.027173893 |
| 2-aminoheptanoate | HMDB0094649 | 0.52 (0.31, 0.87) | 0.0124 | 0.027412818 |
| linoleoyl-docosahexaenoyl-glycerol (18:2/22:6) [2]* | HMDB0007266 | 2.31 (1.20, 4.46) | 0.0124 | 0.027412818 |
| N-acetylmethionine | HMDB0011745 | 2.63 (1.23, 5.65) | 0.0128 | 0.027771247 |
| X-12193 | | 2.02 (1.16, 3.53) | 0.0128 | 0.027771247 |
| X-12456 | | 2.63 (1.22, 5.66) | 0.0132 | 0.028210041 |
| phenylalanine | HMDB0000159 | 1.81 (1.13, 2.89) | 0.0132 | 0.028210041 |
| citrulline | HMDB0000904 | 0.51 (0.30, 0.87) | 0.0133 | 0.028210041 |
| N-acetylthreonine | HMDB0062557 | 2.10 (1.17, 3.78) | 0.0134 | 0.028232134 |
| N1-methyl-2-pyridone-5-carboxamide | HMDB0004193 | 2.13 (1.17, 3.89) | 0.0138 | 0.028833152 |
| 3-amino-2-piperidone | HMDB0000323 | 1.90 (1.13, 3.19) | 0.0147 | 0.030396808 |
| 1-oleoyl-GPE (18:1) | HMDB0011506 | 1.83 (1.13, 2.99) | 0.0147 | 0.030396808 |

| | | | | |
|---|---|---|---|---|
| 1-linolenoyl-GPC (18:3)* | HMDB0010388 | 2.05 (1.15, 3.65) | 0.0149 | 0.030471103 |
| 3-aminoisobutyrate | HMDB0002166 | 0.50 (0.29, 0.88) | 0.0151 | 0.030656814 |
| pro-hydroxy-pro | HMDB0006695 | 3.78 (1.29, 11.08) | 0.0154 | 0.030851217 |
| X-12411 | | 1.97 (1.14, 3.40) | 0.0154 | 0.030851217 |
| phosphate | HMDB0001429 | 2.79 (1.22, 6.41) | 0.0155 | 0.030851217 |
| cerotoylcarnitine (C26)* | HMDB0006347 | 0.57 (0.37, 0.90) | 0.0157 | 0.030851217 |
| asparagine | HMDB0000168 | 0.56 (0.35, 0.90) | 0.0157 | 0.030851217 |
| 3-formylindole | HMDB29737 | 2.13 (1.15, 3.92) | 0.0158 | 0.030878566 |
| quinolinate | HMDB0000232 | 2.30 (1.17, 4.52) | 0.0161 | 0.031259953 |
| indole-3-carboxylate | HMDB0003320 | 2.61 (1.19, 5.74) | 0.0172 | 0.033105505 |
| mannitol/sorbitol | HMDB0000247,HMDB0000765 | 1.89 (1.12, 3.19) | 0.0175 | 0.033586019 |
| lysine | HMDB0003405 | 1.91 (1.12, 3.27) | 0.0180 | 0.034292348 |
| 1-palmitoyl-2-palmitoleoyl-GPC (16:0/16:1)* | HMDB0007969 | 1.85 (1.11, 3.09) | 0.0186 | 0.035125903 |
| isoleucylhydroxyproline* | HMDB0028908 | 2.94 (1.19, 7.23) | 0.0190 | 0.035757853 |
| octadecenedioate (C18:1-DC) | | 0.50 (0.28, 0.89) | 0.0195 | 0.036216555 |
| glycohyocholate | HMDB0000138 | 2.11 (1.13, 3.95) | 0.0196 | 0.036216555 |
| ceramide (d18:2/24:1, d18:1/24:2)* | HMDB0240679,HMDB0240680 | 2.11 (1.13, 3.95) | 0.0197 | 0.036216555 |
| sphingomyelin (d18:2/24:2)* | HMDB0240644 | 1.97 (1.11, 3.49) | 0.0200 | 0.036490668 |
| X-11315 | | 0.55 (0.34, 0.91) | 0.0201 | 0.036490668 |
| N-acetylneuraminate | HMDB0000230 | 2.60 (1.16, 5.84) | 0.0202 | 0.036521325 |
| erythritol | HMDB0002994 | 1.73 (1.09, 2.77) | 0.0212 | 0.037827321 |
| sphingomyelin (d18:1/25:0, d19:0/24:1, d20:1/23:0, d19:1/24:0)* | HMDB0240675,HMDB0240674,HMDB0240673,HMDB0240671 | 1.99 (1.11, 3.58) | 0.0213 | 0.037827321 |
| 3beta-hydroxy-5-cholestenoate | HMDB0012453 | 0.53 (0.31, 0.91) | 0.0213 | 0.037827321 |
| pregnenetriol disulfate* | | 2.27 (1.13, 4.57) | 0.0215 | 0.037953836 |
| X-23665 | | 0.51 (0.29, 0.91) | 0.0217 | 0.038041324 |
| lignoceroylcarnitine (C24)* | HMDB0240665 | 0.56 (0.34, 0.93) | 0.0238 | 0.041462662 |
| 1-palmitoyl-2-arachidonoyl-GPI (16:0/20:4)* | HMDB0009789 | 1.90 (1.09, 3.32) | 0.0239 | 0.041462662 |
| X-11979 | | 2.35 (1.12, 4.95) | 0.0241 | 0.041484703 |
| X-17438 | | 0.50 (0.27, 0.91) | 0.0242 | 0.041484703 |
| 5-hydroxyhexanoate | HMDB0000409,HMDB0000525 | 2.28 (1.11, 4.67) | 0.0251 | 0.042573396 |
| hexadecadienoate (16:2n6) | HMDB0000477 | 1.84 (1.08, 3.12) | 0.0251 | 0.042573396 |
| N4-acetylcytidine | HMDB0005923 | 2.03 (1.09, 3.79) | 0.0257 | 0.043201958 |
| X-23276 | | 1.73 (1.07, 2.81) | 0.0260 | 0.043525874 |
| pregnenediol disulfate (C21H34O8S2)* | | 2.00 (1.08, 3.69) | 0.0264 | 0.043875489 |
| homovanillate (HVA) | HMDB0000118 | 2.06 (1.08, 3.93) | 0.0274 | 0.045005914 |
| cytidine | HMDB0000089 | 2.13 (1.09, 4.16) | 0.0274 | 0.045005914 |
| bilirubin degradation product, C17H20N2O5 (2)** | | 1.90 (1.07, 3.37) | 0.0277 | 0.045005914 |
| 3-hydroxyoctanoylcarnitine (2) | | 0.59 (0.37, 0.94) | 0.0278 | 0.045005914 |
| 5,6-dihydrouridine | HMDB0000497 | 2.07 (1.08, 3.98) | 0.0278 | 0.045005914 |
| arginine | HMDB0000517 | 1.71 (1.06, 2.76) | 0.0287 | 0.046138334 |
| 3-hydroxy-2-ethylpropionate | HMDB0000396 | 1.75 (1.06, 2.90) | 0.0288 | 0.046155386 |
| X-16397 | | 0.57 (0.35, 0.95) | 0.0295 | 0.046958746 |
| p-cresol sulfate | HMDB0011635 | 0.48 (0.25, 0.93) | 0.0302 | 0.047765322 |
| 3-hydroxyoctanoylcarnitine (1) | | 0.59 (0.36, 0.95) | 0.0303 | 0.047765322 |
| vanillylmandelate (VMA) | HMDB0000291 | 2.28 (1.08, 4.80) | 0.0306 | 0.047890994 |
| 1-stearoyl-2-arachidonoyl-GPE (18:0/20:4) | HMDB0009003 | 1.87 (1.06, 3.29) | 0.0316 | 0.049216014 |
| N-formylmethionine | HMDB0001015 | 2.10 (1.07, 4.15) | 0.0319 | 0.049235269 |
| X-26107 | | 0.58 (0.36, 0.96) | 0.0322 | 0.049235269 |

| | | | | |
|---|---|---|---|---|
| 3-hydroxybutyrate (BHBA) | HMDB0000442,HMDB0000357,HMDB0000011 | 0.52 (0.28, 0.95) | 0.0322 | 0.049235269 |
| lactosyl-N-palmitoyl-sphingosine (d18:1/16:0) | HMDB0006750 | 0.52 (0.29, 0.95) | 0.0324 | 0.049235269 |
| 2-oxoarginine* | HMDB0004225 | 1.71 (1.05, 2.80) | 0.0324 | 0.049235269 |
| dihomo-linolenate (20:3n3 or n6) | HMDB0002925 | 1.76 (1.05, 2.97) | 0.0326 | 0.049257472 |
| carnitine | HMDB0000062 | 0.60 (0.38, 0.96) | 0.0333 | 0.049981092 |

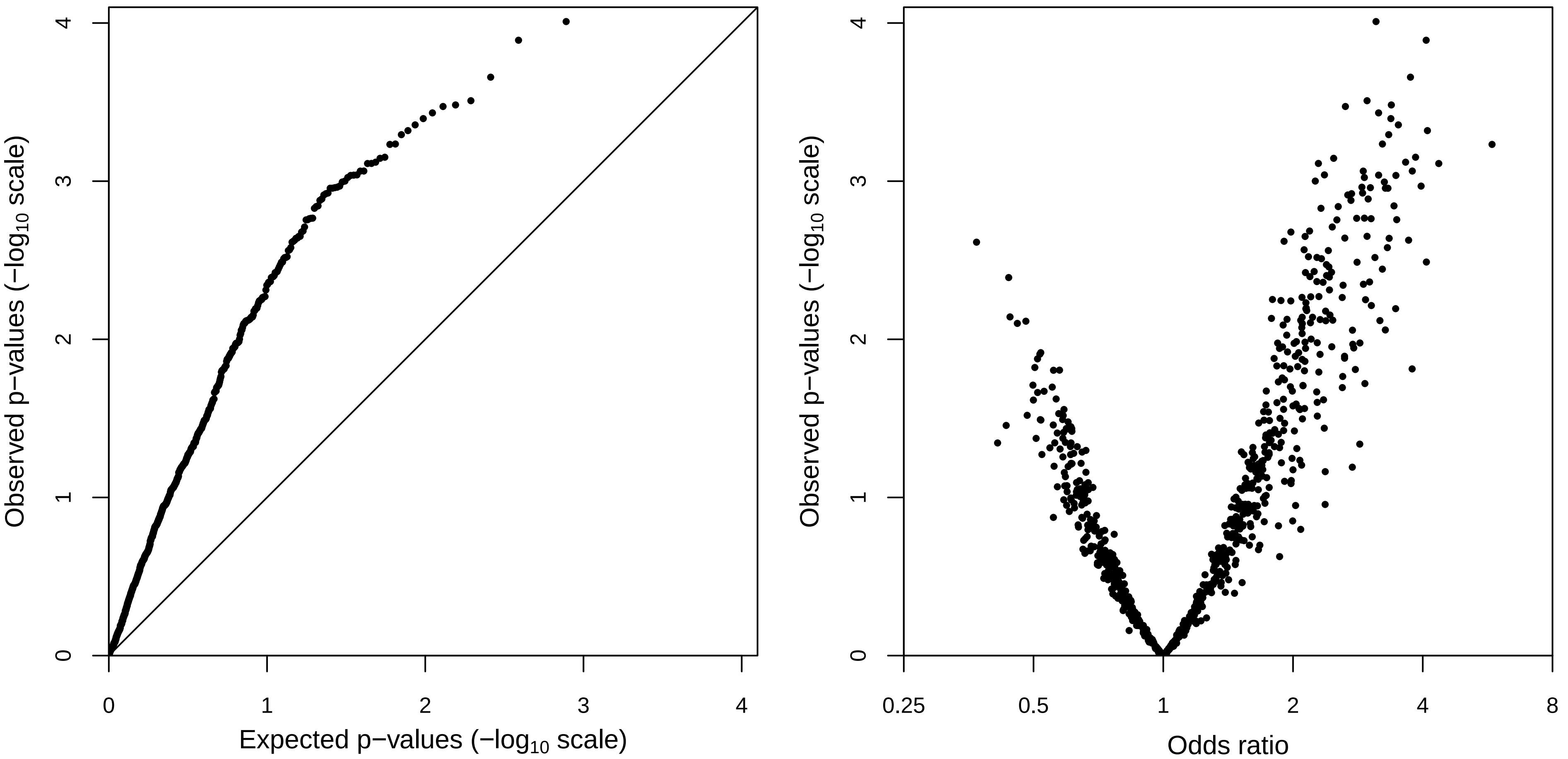

Observed p−values (−log10 scale)
Expected p−values (−log10 scale)
0
1
2
3
4
Observed p−values (−log10 scale)
Odds ratio
0.25
0.5
1
2
4
8

Supplementary Table 2. Clinical variables and metabolites selected by elastic net regression analysis

| Variable | Clinical data only | Metabolites only | Metabolites + clinical data |
|---|---|---|---|
| Age | - | | |
| Sex | + | | |
| History of thrombosis | + | | + |
| STAT score | + | | + |
| N6-methyladenosine | | - | - |
| Aspartate | | | |
| Cortisone | | - | - |
| Sphingosine | | + | + |
| Isoleucine | | | + |
| Glucose | | | |
| Pyruvate | | + | + |
| Thymidine | | + | + |
| Dihydroorotate | | | |
| Caprylate (8:0) | | | |
| Pipecolate | | | |
| Phosphoethanolamine | | + | |
| Gamma-glutamyltyrosine | | | |
| Gamma-glutamylleucine | | | |
| N-stearoyl-sphingosine (d18:1/18:0)* | | + | |
| Eicosapentaenoate (EPA; 20:5n3) | | | |
| Carnitine | | | + |
| Glycerol 3-phosphate | | | |
| 3-indoxyl sulfate | | | |
| Gamma-glutamylphenylalanine | | | |
| 1-stearoyl-2-oleoyl-GPS (18:0/18:1) | | | |
| Erythritol | | + | + |
| 1-oleoylglycerol (18:1) | | + | + |
| 2-linoleoylglycerol (18:2) | | | |
| 1-linoleoylglycerol (18:2) | | + | + |
| Propionylcarnitine (C3) | | | |
| 3-carboxy-4-methyl-5-propyl-2-furanpropanoate (CMPF) | | | |
| N-acetylphenylalanine | | | |
| Gamma-glutamylthreonine | | | |
| P-cresol sulfate | | + | |
| N6-carbamoylthreonyladenosine | | | + |
| 3-methylcytidine | | - | - |
| N1-methyl-2-pyridone-5-carboxamide | | | |
| Docosapentaenoate (n6 DPA; 22:5n6) | | | |
| Mannitol/sorbitol | | | |
| Thymol sulfate | | | |
| Sphingomyelin (d18:1/18:1, d18:2/18:0) | | | |
| Andro steroid monosulfate C19H28O6S (1)* | | | |
| 2R,3R-dihydroxybutyrate | | | |
| 2,3-dihydroxyisovalerate | | | |
| Formiminoglutamate | | | |
| Glycohyocholate | | | |
| Guaiacol sulfate | | | |
| 2-aminoheptanoate | | | + |
| Methyl-4-hydroxybenzoate sulfate | | + | + |
| N-acetylkynurenine (2) | | | |
| 1-stearoyl-2-arachidonoyl-GPE (18:0/20:4) | | | + |
| 1-(1-enyl-palmitoyl)-2-linoleoyl-GPC (P-16:0/18:2)* | | | |
| 1-linoleoyl-GPG (18:2)* | | | |
| Ceramide (d18:2/24:1, d18:1/24:2)* | | | |
| 2-naphthol sulfate | | | |
| Deoxycholic acid glucuronide | | | + |
| 1-methyl-5-imidazolelactate | | + | |
| X-15503 | | | |
| X-15674 | | | |
| X-17438 | | | |
| X-22162 | | | |
| X-23276 | | | |
| X-23665 | | | |
| X-24541 | | | + |
| X-25267 | | | |

X: indicating unnamed metabolites.
+: indicating a positive regression coefficient for the metabolite; -: indicating a negative regression coefficient for the metabolite.
